# An integrative approach identifies direct targets of the late viral transcription complex and an expanded promoter recognition motif in Kaposi’s sarcoma-associated herpesvirus

**DOI:** 10.1101/550772

**Authors:** Divya Nandakumar, Britt Glaunsinger

## Abstract

The structural proteins of DNA viruses are generally encoded by late genes, whose expression relies on recruitment of the host transcriptional machinery only after the onset of viral genome replication. β and γ-herpesviruses encode a unique six-member viral pre-initiation complex (vPIC) for this purpose, although how the vPIC directs specific activation of late genes remains largely unknown. The specificity underlying late transcription is particularly notable given that late gene promoters are unusually small, with a modified TATA-box being the only recognizable element. Here, we explored the basis for this specificity using an integrative approach to evaluate vPIC-dependent gene expression combined with promoter occupancy during Kaposi’s sarcoma-associated herpesvirus (KSHV) infection. This approach distinguished the direct and indirect targets of the vPIC, ultimately revealing a novel promoter motif critical for KSHV vPIC binding. Additionally, we found that the KSHV vPIC component ORF24 is required for efficient viral DNA replication and identified a ORF24 binding element in the origin of replication that is necessary for late gene promoter activation. Together, these results identify an elusive element that contributes to vPIC specificity and suggest novel links between KSHV DNA replication and late transcription.

**Author summary:** Gene expression in DNA viruses often occurs in temporal waves, with expression of essential structural proteins occurring late in infection, after viral genome replication has begun. Strategies underlying expression of these viral late genes are often sophisticated; for example, the β- and γ-herpesviruses encode a six-component viral complex that directs late gene transcription, largely by unknown mechanisms. Here, we evaluated how this complex specifically recognizes late promoters during infection with the oncogenic human γ-herpesvirus Kaposi’s sarcoma-associated herpesvirus (KSHV). We found that one of the components of the late transcription complex was required for robust viral DNA replication and binds at the origin of replication, suggesting new links between KSHV replication and transcription. Combined measurements of late gene expression and promoter occupancy then revealed which KHSV genes are directly controlled by the late gene transcription complex, leading to identification of a key new regulatory element in KSHV late promoters. Together, these data help explain how the late gene transcription complex is able to bind seemingly minimal promoters with high specificity, ensuring robust expression of viral factors necessary for assembly of progeny virions.

## Introduction

A conserved feature of double-stranded DNA viruses is that they possess a set of genes that become strongly activated only following viral DNA replication (1). These “late genes” generally encode factors involved in viral assembly that are essential for virion production. The mechanisms underlying their transcriptional regulation are often non-canonical or distinct from those involved in activating either cellular genes or early viral genes. For example, baculoviruses encode a multisubunit, α-amanitin-resistant DNA-directed RNA polymerase dedicated specifically to the activation of compact late promoters (2). Similarly, bacteriophage T7 encodes its own late gene specific RNA polymerase that recognizes unique promoter elements to transcribe late genes (3,4). Adenoviruses regulate late gene expression using a complex combination of viral and cellular factors; beyond promoter activation, these also regulate transcription termination and contribute to differential posttranscriptional processing of late viral mRNA (5).

Several studies suggest that the expression of late genes is also uniquely regulated in β- and γ-herpesviruses, which include human cytomegalovirus (HCMV), Epstein-Barr virus (EBV), and Kaposi’s sarcoma-associated herpesvirus (KSHV) (6). Unlike other eukaryotic dsDNA viruses, β and γ herpesviruses do not engage cellular TATA binding protein (TBP) during late gene pre-initiation complex formation (7). Their core late gene promoters are unusually small (12-15 base pairs (bp)), with the only recognizable conserved element being a modified TATA box (TATT) ~30 bp upstream of the transcription start site (TSS) (8–10). This is in contrast to eukaryotic and most early herpesviral promoters, which have multiple promoter elements that are important for promoter recognition and activation by RNA polymerase II (RNA pol II) (11). Another notable feature is that activation of these minimal promoters requires a virally-encoded six-protein complex called the viral Pre-Initiation Complex (vPIC) that coordinates mammalian RNA pol II (12,13). The vPIC is presumed to be uniquely adapted to activate minimal promoters, yet the mechanisms underlying its activity and specificity remain largely unknown.

The core component of the KSHV vPIC is ORF24 (BcRF1 in EBV and UL87 in HCMV), a viral TBP mimic that displays putative structural similarity to cellular TBP (7,14,15). Unlike cellular TBP, the viral TBP mimics have expanded functionality in that they bind RNA pol II in addition to promoter DNA, making them unique among transcription factors that function in eukaryotic cells. Beyond ORF24, the other essential components of the KSHV vPIC are ORFs 18, 30, 31, 34, and 66, but little is known about how each of these factors contributes to vPIC activity (7,16–18). ORF34 interacts with multiple components of the vPIC, and is thus thought to function as a scaffolding factor (13). The ORF18 homolog in HCMV (pUL79) has been shown to function as an elongation factor, although whether it plays a similar role in KSHV and EBV is unclear (19). Despite the paucity of functional information on the individual vPIC components, protein-protein interaction mapping has provided insight into the overall complex organization (13,18). Furthermore, point mutants that disrupt individual inter-subunit vPIC interactions abrogate late gene transcription, highlighting the fact that vPIC complex integrity is critical for function (13,20,21).

The mechanistic basis for promoter selectivity has also been challenging to grasp in the context of the minimal, largely nondescript nature of late gene promoters. While a modified TATA box with a TATT sequence is enriched in several late promoters of KSHV, MHV68, and EBV (8– 10,22), the ORF24 TBP mimic shows partial binding to a mutant promoter in which the TATT sequence is modified to resemble a canonical TATA box (8). However, ORF24 does not bind promoters that inherently have a canonical TATA box (8), which suggests that other DNA elements surrounding the TATT sequence in late gene promoters contribute to ORF24 binding or vPIC assembly on the DNA.

One possible reason that additional DNA elements involved in vPIC promoter recognition have not been identified is that we do not yet know the subset of viral promoters under the direct influence of the vPIC. Techniques such as RNA Sequencing (RNA Seq) and microarrays have helped define the set of genes that are expressed with late kinetics and in a manner dependent on viral DNA replication (23–26). Furthermore, viral mutagenesis and knockdown studies have revealed viral genes whose expression is impaired in the absence of vPIC components (17,18,22,27,28). However, these assays cannot distinguish direct vPIC targets from indirect targets, such as genes regulated by factors that are themselves dependent on the vPIC. Inclusion of such indirect targets in the pool of vPIC-regulated promoters could confound the identification of conserved elements important for direct vPIC regulation.

Here, we integrated RNA Seq with chromatin immunoprecipitation high-throughput sequencing (ChIP Seq) for vPIC components during KSHV infection to distinguish genes whose expression is directly versus indirectly controlled by the vPIC. In doing so we made four key findings that provide insight into late gene promoter activation in KSHV. First, we defined the set of direct vPIC promoter targets within the KSHV genome and showed that other promoters are indeed nonproductively bound or only indirectly regulated by the vPIC. Second, by restricting our analysis to these direct targets, we revealed a novel promoter motif that contributes significantly to vPIC binding and thus provides insight into features governing vPIC specificity. Notably, this motif did not emerge when the indirect vPIC targets were included in the analysis. Third, we found that the KSHV TBP mimic ORF24 is required for efficient viral DNA replication. This was unexpected given that the other studied vPIC components have been shown to be dispensable for this process. Finally, we identified a binding site for ORF24-34 at the lytic origin of replication that is essential for activating late gene expression, providing a new connection between viral DNA replication and late gene transcription.

## Results

### KSHV ORF24 is essential for late gene expression and contributes to viral DNA replication

Three leucine residues in the N-terminal domain of KSHV ORF24 (amino acids 73-75) are essential for its interaction with RNA Pol II and its ability to promote late gene transcription (7). Given that most viral proteins are multifunctional, we engineered the ORF24_RLLLG->RAAAG_ mutation (ORF24_RAAAG_) into KSHV BAC16 to precisely evaluate the impact of the transcriptional function of ORF24 during lytic infection. We also generated the corresponding mutant rescue (ORF24_RAAAG_-MR) to ensure that any observed phenotypes were not the effect of secondary mutations in the BAC. Identical banding patterns from *Rsr*II digestion of WT and mutant BAC16 DNAs verified that there were no gross rearrangements of the viral genome during mutagenesis (Figure S1-lanes1-3). We established stable cell lines latently infected with WT, ORF24_RAAAG_, and ORF24_RAAAG_-MR KSHV in the iSLK background, which contains a doxycycline-inducible version of the KSHV lytic transactivator protein ORF50 (RTA) to enable lytic reactivation (29). During lytic reactivation, these three viruses expressed similar levels of the early protein ORF59, while the ORF24_RAAAG_ mutant was selectively unable to express the K8.1 late protein or produce infectious virions (Fig 1A, 1B). These findings are consistent with previous complementation data (7), and confirm the importance of the ORF24-Pol II interaction for late gene expression.

**Fig 1:**
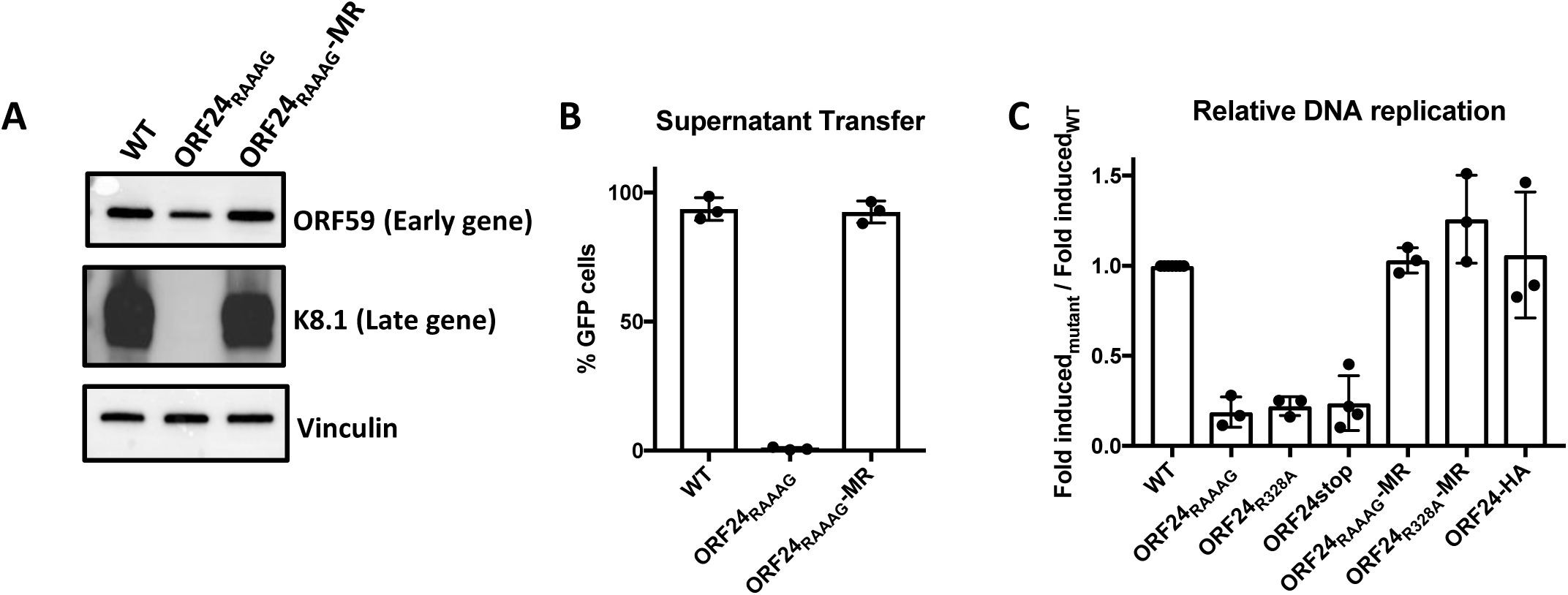
The ORF24-RNA pol II interaction is essential for late gene expression and contributes to viral DNA replication. iSLK cells harboring WT BAC16, the ORF24_RAAAG_ mutant or the ORF24_RAAAG_ mutant rescue (MR) were reactivated with doxycycline and sodium butyrate for 72 hours. (A) Western blot showing expression of an early protein (ORF59) and a late protein (K8.1) in WT, ORF24_RAAAG_ and ORF24_RAAAG_-MR. Vinculin was used as a loading control. (B) Progeny virion production was measured by supernatant transfer on to 293T cells and quantified by flow cytometry. (C) DNA replication of different mutants and MRs was measured by qPCR and normalized to the respective uninduced sample. The fold induction of all mutants was normalized to the WT to get the relative change in DNA replication compared to WT.

Surprisingly, we also observed ~6-fold defect in DNA replication in the ORF24_RAAAG_ mutant (Fig 1C), which was unexpected given that previous studies with mutations in ORF24 or other components of the vPIC did not show this defect (7,13,16–18). This defect was rescued in the ORF24_RAAAG_-MR cell line, indicating that it was not due to secondary mutations in the BAC (Fig. 1C). However, to independently confirm this observation, we engineered two additional ORF24 viral mutants: the point mutant ORF24_R328A_, which abrogates its essential interaction with the ORF34 component of the vPIC (13), and a stop mutant (ORF24_Stop_) that has a premature stop codon in the ORF24 coding sequence. Inclusion of the ORF24_Stop_ virus allowed us to evaluate whether the point mutants were functioning in a dominant negative manner, for example through nonproductive DNA binding. Finally, we also N-terminally tagged endogenous WT ORF24 with HA (HA-ORF24) to assess whether any alterations to this locus impacted viral DNA replication. Notably, we observed a similar ~5-6-fold defect in DNA replication in the ORF24_RAAAG_, ORF24_R328A_, and ORF24_Stop_ viruses, which was restored to WT levels in the ORF24_RAAAG_ and ORF24_R328A_ MR viruses. Moreover, there was no DNA replication defect in the HA-ORF24 virus (Fig 1C), demonstrating that the defect in DNA replication is not a consequence of manipulating the ORF24 locus but is specific to disrupting its role in transcription.

### Specific gene clusters show differential dependence on the ORF24-Pol II interaction

To more comprehensively analyze how the ORF24-Pol II interaction impacts gene expression during infection, we deep sequenced RNA from three replicates each of iSLK cells infected with the ORF24_RAAAG_ or the ORF24_RAAAG_-MR virus. Samples were sequenced at 24 and 48 h post lytic reactivation with doxycycline and sodium butyrate, as these times captured predominantly early (24 h) versus both early and late (48 h) viral gene expression (Figure S2). Given that overlapping transcripts are pervasive in the KSHV genome (22,30), we calculated the primary transcript counts as described in Bruce et al (30) to accurately quantify transcript levels.

Differential expression analysis of genes in infected cells containing WT ORF24 (ORF24_RAAAG_ MR) confirmed that the majority of viral genes are upregulated between 24 and 48 h post reactivation (Figure 2A), and those that are most strongly upregulated are predominantly classified as late genes (shown in blue). Conversely, there was a generalized reduction in transcript levels at the 48 h time point in cells containing KSHV ORF24_RAAAG_ compared to MR, with the most strongly downregulated genes being those classified as late genes (Figure 2B).

**Fig 2:**
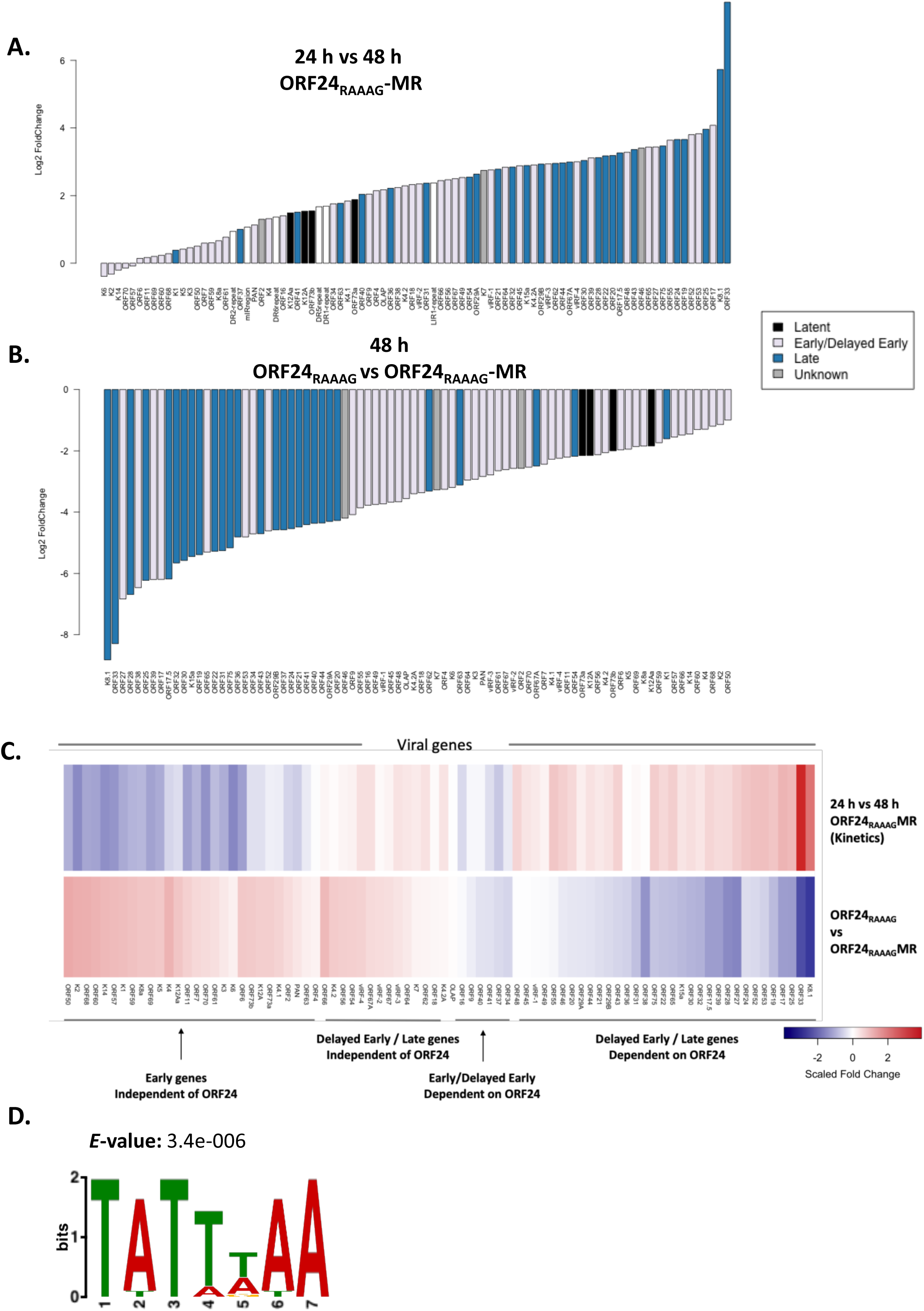
Differential expression analysis of ORF24-Pol II interaction mutant and mutant rescue. iSLK cells harboring ORF24_RAAAG_ and ORF24_RAAAG_-MR were reactivated with doxycycline and sodium butyrate for 24 and 48 h, whereupon total RNA was extracted and subjected to library preparation and RNA sequencing. (A) Bar plot showing log_2_ fold change of the viral genes in the ORF24_RAAAG_-MR at 48 h relative to 24 h. Genes were color coded by kinetic class as defined in (30). (B) Bar plot showing log_2_ fold change of the viral genes in the mutant relative to the MR. Genes were color coded by kinetic class as defined in (30). (C) Heatmap of the scaled log_2_ fold change of the genes induced in the MR between 24 to 48 h and genes repressed in the mutant compared to MR at 48 h. Gene clusters with different kinetics and dependence on ORF24-Pol II interaction are indicated. (D) Consensus motif identified by MEME analysis of promoters from 14 genes that showed greater than one standard deviation repression in the mutant relative to the MR.

**Fig 3:**
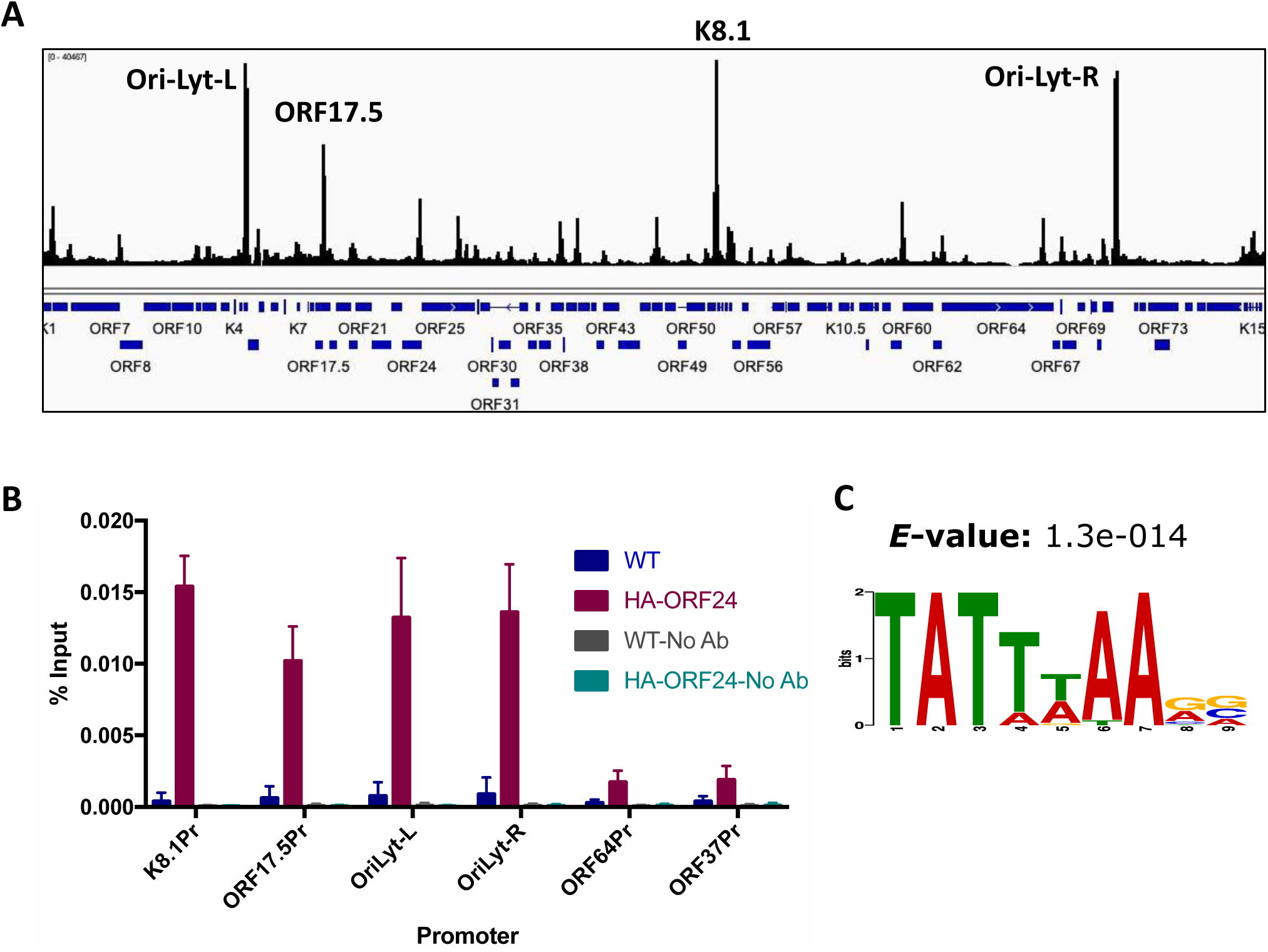
ChIP-Seq with HA-ORF34 shows multiple binding sites across the viral genome. (A) ChIP-Seq was performed using an anti-HA antibody to immunoprecipitate DNA prepared from iSLK HA-ORF34 cells reactivated for 48 h with doxycycline and sodium butyrate. iSLK-WT cells lacking the HA tag and input DNA were used as negative controls. The alignment files were converted to the tiled data file (tdf) format and the ChIP background from the untagged cells was subtracted and visualized on the Integrated Genome Viewer (IGV) (B) ChIP-qPCR was performed using an anti-HA antibody to immunoprecipitate DNA prepared from iSLK HA-ORF24 cells reactivated for 48 h with doxycycline and sodium butyrate. The DNA was quantified by qPCR with primers specific to the promoter regions of the indicated genes. Data were normalized to the level of input DNA and are presented as percentages of the input. Data shown are from three biological replicates. (C) Consensus motif identified by MEME analysis of promoters from 19 genes whose promoters had a ChIP peak.

Combining the above ORF24_RAAAG_ mutant and MR induction data into a heatmap confirmed that in general, genes with later expression kinetics tended to be more severely impacted by the ORF24_RAAAG_ mutation compared to genes with early expression kinetics (Figure 2C). (The fold change data was scaled to allow for easy visualization). However, not all genes fit this model. Some genes categorized as early based on their induction pattern showed a moderate dependence on the ORF24-Pol II interaction (e.g. ORFs 9, 40, 34, 41, 16 and 37). Similarly, a subset of genes that show late kinetics were less dependent on the ORF24-Pol II interaction (e.g. ORFs 67A, 62, K4.2A, 64, K7, 54, 67, 56, 66, K4.2 and vIRFs-2, 3 and 4). Classification as early or late and dependence on the ORF24-Pol II interaction was determined using mean log_2_ fold change. Together, these data suggest that not all kinetic late genes are regulated by the vPIC and there may be subset of early genes that are direct or indirect targets of the vPIC.

### The TATTWAA motif is enriched in the promoters of genes whose expression is linked to the ORF24-Pol II interaction

We then looked for motifs enriched in the promoters of genes most strongly downregulated in the ORF24_RAAAG_ mutant using Multiple EM for Motif Elicitation (MEME) (31), a program that discovers un-gapped motifs that are enriched in a given set of sequences. We defined the promoter sequence as 100 bp upstream from the transcription start site (TSS) or 300 bp upstream from the translation start site when the TSS was unknown. Analysis of promoter sequences from 14 of the most strongly down-regulated genes (log_2_ fold change greater than one standard deviation from the mean) showed that TATTWAA is the only motif that is enriched in this set of sequences (Figure 2D).

We next monitored for the occurrence of this motif across all promoters in the viral genome using Find Individual Motif Occurrences (FIMO) (32), which scans a set of sequences for matches to motifs provided by the user. 23 of 83 viral promoters were found to have the TATTWAA motif with a high score, of which 16 promoters had the motif ~30 bp upstream from the TSS (Figure 4 and Table 1), consistent with the location of the canonical TATA box in eukaryotic promoters. Of the remaining seven promoter sequences, the TSS of six of the promoters is not known (shown in red in Table 1); hence, the exact position of the TATTWAA motif from the TSS cannot be estimated. ORF25 is the only exception, where the TATTWAA sequence is not present ~30 bp upstream from the TSS, although it does have a canonical TATA box (TATAAAA) at that position.

**Table 1.** Occurrence of TATTWAA motif in KSHV

| Promoter | Start | Stop | Strand | Score | p-value | q-value | Matched Sequence | Position |
| --- | --- | --- | --- | --- | --- | --- | --- | --- |
| ORF17 | 68 | 74 | + | 11.8614 | 0.000114 | 0.143 | TATTTAA | 32 |
| ORF62 | 68 | 74 | + | 11.8614 | 0.000114 | 0.143 | TATTTAA | 32 |
| ORF55 | 69 | 75 | + | 11.8614 | 0.000114 | 0.143 | TATTTAA | 31 |
| ORF39 | 70 | 76 | + | 11.8614 | 0.000114 | 0.143 | TATTTAA | 30 |
| ORF8 | 71 | 77 | + | 11.8614 | 0.000114 | 0.143 | TATTTAA | 29 |
| ORF52 | 71 | 77 | + | 11.8614 | 0.000114 | 0.143 | TATTTAA | 29 |
| ORF17.5 | 71 | 77 | + | 11.8614 | 0.000114 | 0.143 | TATTTAA | 29 |
| ORF59 | 71 | 77 | + | 11.8614 | 0.000114 | 0.143 | TATTTAA | 29 |
| ORF58 | 71 | 77 | + | 11.8614 | 0.000114 | 0.143 | TATTTAA | 29 |
| ORF19 | 187 | 193 | + | 11.8614 | 0.000114 | 0.143 | TATTTAA | 113 |
| ORF27 | 197 | 203 | + | 11.8614 | 0.000114 | 0.143 | TATTTAA | 103 |
| ORF32 | 243 | 249 | + | 11.8614 | 0.000114 | 0.143 | TATTTAA | 57 |
| ORF25 | 11 | 17 | + | 11.198 | 0.000235 | 0.153 | TATTAAA | 89 |
| K8.1 | 68 | 74 | + | 11.198 | 0.000235 | 0.153 | TATTAAA | 32 |
| K4.1 | 68 | 74 | + | 11.198 | 0.000235 | 0.153 | TATTAAA | 32 |
| ORF65 | 69 | 75 | + | 11.198 | 0.000235 | 0.153 | TATTAAA | 31 |
| ORF33 | 70 | 76 | + | 11.198 | 0.000235 | 0.153 | TATTAAA | 30 |
| ORF26 | 71 | 77 | + | 11.198 | 0.000235 | 0.153 | TATTAAA | 29 |
| ORF38 | 71 | 77 | + | 11.198 | 0.000235 | 0.153 | TATTAAA | 29 |
| ORF4 | 71 | 77 | + | 11.198 | 0.000235 | 0.153 | TATTAAA | 29 |
| ORF64 | 145 | 151 | + | 11.198 | 0.000235 | 0.153 | TATTAAA | 155 |
| ORF66 | 218 | 224 | + | 11.198 | 0.000235 | 0.153 | TATTAAA | 82 |
| ORF22 | 229 | 235 | + | 11.198 | 0.000235 | 0.153 | TATTAAA | 71 |

ORFs shown in red do not have a known Transcription Start Site (TSS)

**Fig 4:**
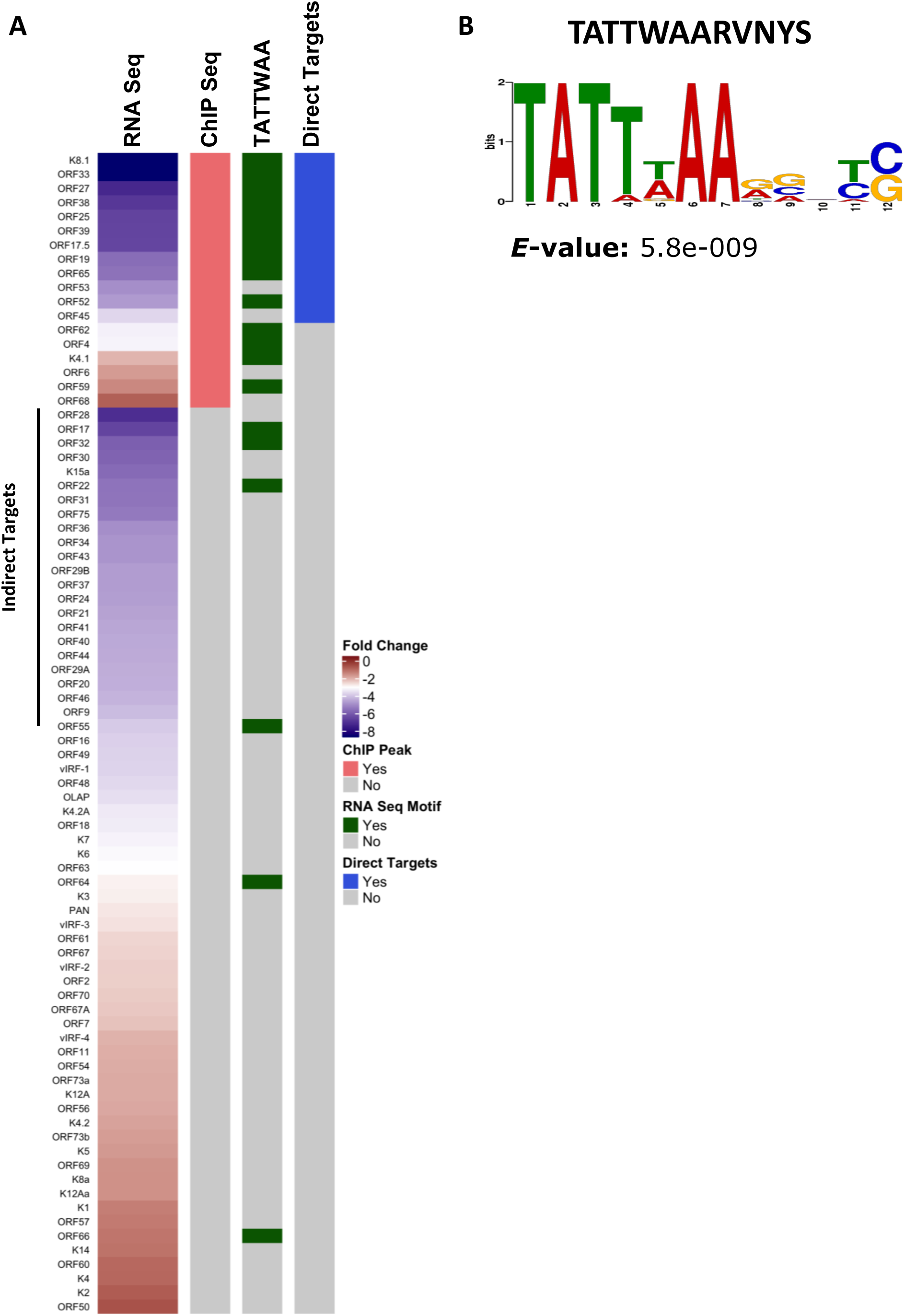
Identifying direct targets of the vPIC. (A) Heatmap showing genes repressed in the ORF24_RAAAG_ mutant compared to the MR adjacent to the list of genes that show ORF34 binding at their promoter by ChIP-Seq analysis. Also indicated are viral genes that were identified to have the TATTWAA motif by FIMO analysis. The direct targets were identified by their combined dependence on the ORF24-Pol II interaction and the presence of a ChIP signal at the promoter. (B) Consensus motif identified by MEME analysis of promoters from 13 direct targets.

Notably, a subset of the genes containing the TATTWAA motif showed less than average repression by the ORF24_RAAAG_ mutation (ORF55, ORF64, ORF66, ORF59, K4.1, ORF4, ORF8, ORF58, ORF26 and ORF62). One explanation for this observation is that these genes could be regulated by cellular transcription factors and other *cis* elements which are absent in the genes that are downregulated in the ORF24_RAAAG_ mutant. The data also suggest that the motif may not be sufficient to mark regulation by the vPIC, which is consistent with a similar observation in MHV68 (8). Furthermore, several genes (ORF28, ORF30, K15, ORF31 and ORF75) were strongly affected by the mutation (log_2_ fold change greater than one standard deviation from the mean) but lack the TATTWAA motif. The number of genes in this set increases to 26 if we broaden the criteria to include all the genes that show greater than average repression in the presence of the mutation. These genes could either be down regulated by an indirect mechanism in the absence of the ORF24-Pol II interaction or could be bound by the vPIC in a manner independent of the TATT motif.

### Chromatin immunoprecipitation reveals vPIC occupancy across the KSHV genome

Our RNA Seq experiments identified KSHV genes whose expression is influenced by the transcription regulatory function of ORF24. However, RNA Seq does not distinguish genes that are directly bound by the vPIC from those that are indirect transcriptional targets (e.g. controlled by other factors whose expression is induced by ORF24). We therefore sought to identify promoters bound by the KSHV vPIC complex using ChIP Seq. Previous studies suggest that ORF34 acts as a scaffolding factor that brings together all components of the vPIC (13) including ORF24, which binds both RNA Pol II and TATT-containing DNA. We reasoned that ORF34 promoter occupancy would therefore be a good proxy for vPIC assembly, and additionally would avoid potential pitfalls associated with any non-productive or independent interactions between ORF24 and DNA.

To facilitate ChIP Seq, we added an N-terminal HA tag to endogenous ORF34 in KSHV BAC16 (HA-ORF34) (Figure S3A), and generated a stable, doxycycline-inducible iSLK cell line containing this tagged virus as previously described. HA-ORF34 produced WT levels of infectious virions in a supernatant transfer assay (Figure S3B), indicating that the tag does not disrupt ORF34 activity. The integrity of the BAC DNA was verified by *Rsr*II digestion (Figure S1-Lane7). We performed the ChIP Seq experiment in duplicate at 48 h post reactivation to ensure active late gene expression. Reactivated iSLK cells with untagged BAC16 served as a negative control for the immunoprecipitation. The individual ChIP peaks for the two replicates as well as the respective inputs and controls are shown in Figure S3C and supplementary file 1. The ChIP Seq data showed that HA-ORF34 bound several locations across the viral genome (Figure 3A) but was not detectably associated with the host genome. A peak calling algorithm (HOMER) (33) identified 25 common peaks in the viral genome across two replicates that were significantly enriched in HA-ORF34 relative to untagged WT or input DNA (Supplementary File 1 and Figure S3C). Notably, 20 of the 25 peaks were found upstream of ORFs, with ~80% of the peaks centered less than 200 bp from the closest TSS (Supplementary File 1). Given the link between DNA replication and late transcription in herpesviruses, it was particularly interesting that two of the most prominent peaks are upstream of the two minimal lytic origins of replication (Ori) in the KSHV genome (Figure 3A). One of the peaks could not be uniquely assigned to a single ORF and was not considered further. Of the remaining 19 ORFs associated with peaks, 13 are involved in packaging, capsid assembly (or form part of capsid itself), 4 are involved in immune evasion and 2 in DNA replication (Table 2). While several of them have also been previously classified as late genes (12/20), it is notable that 6 of them have been classified as early genes, again suggesting that not all targets of the vPIC are late genes.

**Table 2.**
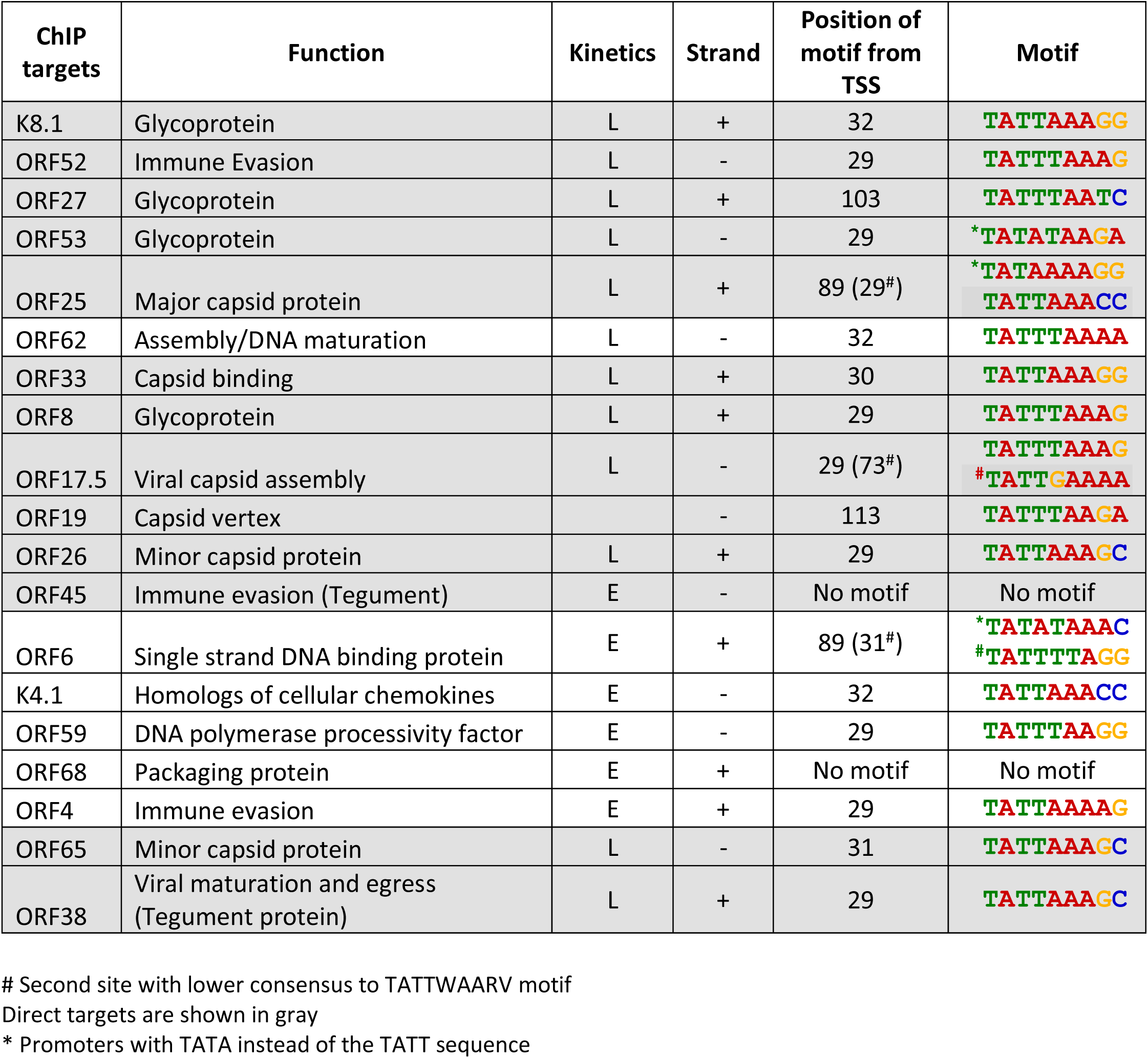
Features of ChIP-Seq targets

To confirm that the observed binding pattern was representative of vPIC assembly, we independently validated several prominent ChIP Seq peaks by ChIP-qPCR from cells infected with KSHV containing HA-tagged ORF24 (HA-ORF24). Similar to HA-ORF34, the HA-ORF24 virus produced WT levels of infectious virions (Figure S3B). Indeed, the HA-ORF24 ChIP-qPCR data were consistent with the HA-ORF34 ChIP-Seq data (Figure 3B), with peaks showing higher percent input values compared to control regions (ORF64 and ORF37 promoters).

A MEME analysis of the promoters associated with the peaks again showed that the only motif enriched in these promoters is TATTWAARV (Figure 3C). 15 of the 19 promoters have the TATTWAA motif and in a majority of the sequences, it is present ~30 bp from the TSS (Table 2). Of the four exceptions, two of the promoters (ORF53 and ORF6) have a TATATAA motif with an A at position 4 in place of the T. Indeed, mORF53 has previously shown to be a target of the vPIC in MHV68 despite the deviation from consensus (8). Two key points can be drawn from the ChIP seq data. First, several genes whose promoters contain a TATTWAA motif do not show ORF34 binding (e.g. ORFs 17, 32, 22, 55, 58, 64 and 66). Second, while most promoters bound by ORF34 have a TATTWAA motif, there are some exceptions (ORF45, ORF68, ORF6 and ORF53). These observations indicate that while the TATTWAA motif is enriched in promoters regulated by the vPIC, it is not sufficient to ensure binding of this specialized transcription complex.

### Thirteen KSHV genes are direct targets of the vPIC

Combining the RNA Seq and ChIP Seq data, we defined the direct targets of the vPIC as genes that are both downregulated in the absence of the ORF24-Pol II interaction and show ORF34 binding in the promoter region. By this measure, a total of 13 genes are direct targets of the vPIC (Figure 4 and Table 2 – gray rows). Of the 13 direct targets, 11 code for structural proteins or proteins involved in viral egress while two are tegument proteins involved in immune evasion. Genes whose expression was impaired in the KSHV ORF24_RAAAG_ mutant but lacked ORF34 binding at their promoters are likely to be indirect targets of the vPIC (Figure 4A). Finally, it is interesting to note that not all genes that have a ChIP peak are down regulated in the RNA Seq data (e.g. ORFs 4, 6, K4.1, 59, 62 and 68). These could be false positive ChIP peaks and/or have alternate mechanisms of regulation that ensure their transcription even in the absence of the vPIC.

### The direct vPIC targets reveal an expanded promoter motif

A defining feature of characterized late gene promoters is that they are minimalistic, with no conserved motifs beyond the TATTWAA signature recognized by ORF24. Yet, our data indicate that the TATTWAA motif is insufficient to confer activation by the late gene transcription complex. We hypothesized that the inclusion of potentially indirect vPIC targets into prior motif searches may have hindered identification of other regulatory sequences within these promoters. Indeed, repeating the MEME analysis on the 13 direct targets revealed that an additional 5 bp motif (RVNYS) downstream of the TATTWAA motif was enriched in these promoter sequences (Figure 4B).

To explore the contribution of the 5 bp region downstream of the TATTWAA sequence for regulation by the vPIC, we used a plasmid-based promoter activation assay in which expression of a luciferase reporter was driven by either the K8.1 late or ORF57 early promoter (Figure 5A). The KSHV left lytic origin of replication (ori-lyt) was included on the plasmids, as data from MHV68, KSHV and EBV indicate that the presence of ori-lyt in *cis* significantly boosts late gene promoter activation during infection (8,10,34). Each plasmid was transfected together with a plasmid constitutively expressing renilla luciferase (to normalize for transfection efficiency) into lytically reactivated KSHV-positive iSLK cells and the luciferase signal was measured 48 h post transfection and reactivation. As expected, the presence of ori-lyt enhanced K8.1 promoter-driven luciferase expression by 15 – 20-fold (Figure 5B). Furthermore, unlike the consistently robust ORF57 promoter activity, K8.1 promoter activity on the plasmid was completely repressed in the presence of DNA replication inhibitor PAA (Figure 5C) or in cells lytically infected with the ORF24_RAAAG_ mutant (Figure 5D). Taken together, these data confirm that the plasmid assay accurately recapitulates late gene promoter activity.

**Fig 5:**
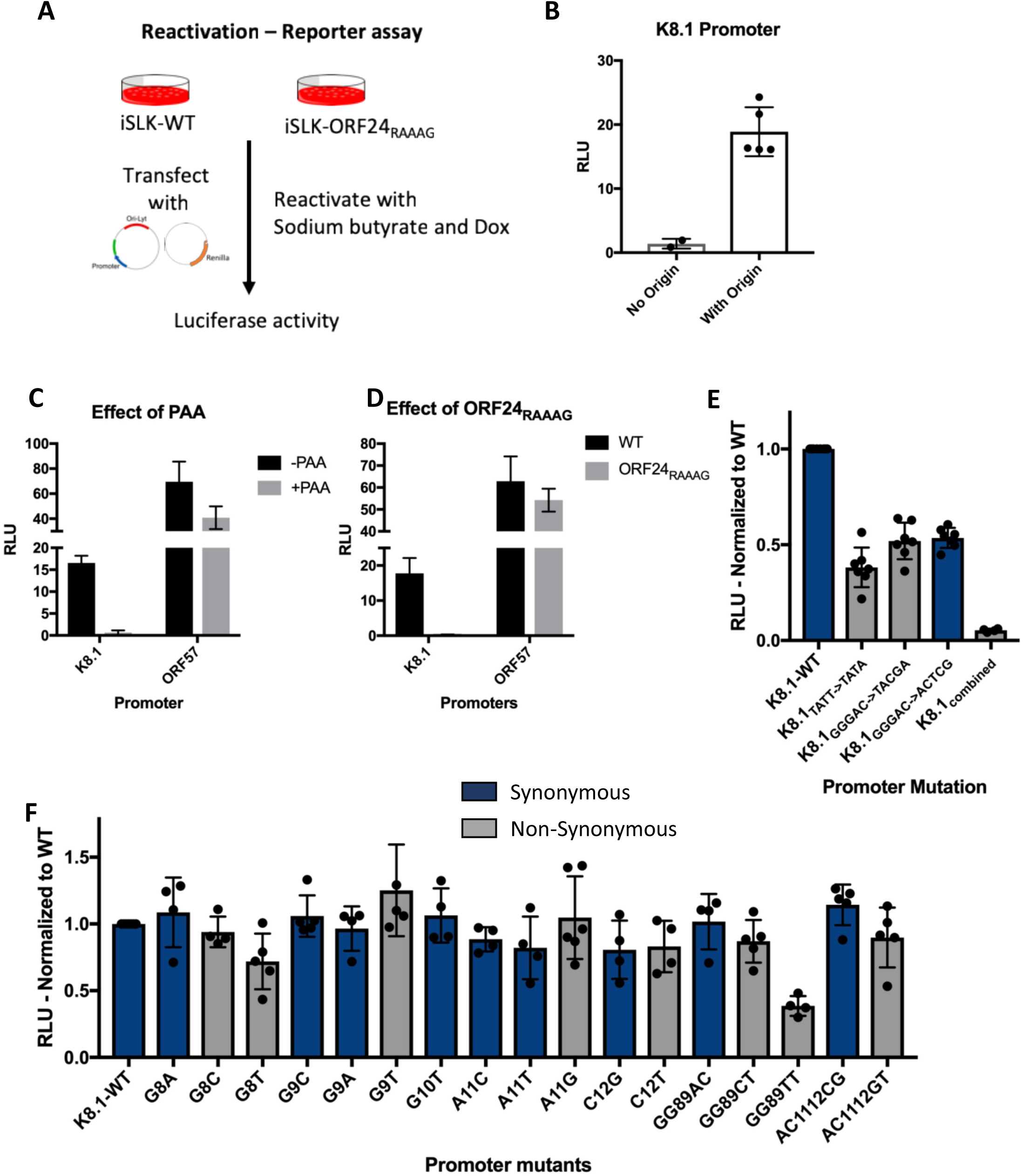
The motif identified from direct targets is important for transcription from the promoter. (A) Schematic of plasmid-based promoter activation assay. iSLK-WT or ORF24_RAAAG_ mutant cells reactivated with doxycycline and sodium butyrate were transfected with reporter plasmid carrying the indicated promoter sequence upstream of the firefly luciferase gene and a control renilla luciferase expressing plasmid. 48 h post transfection and reactivation, the cells were assessed for luciferase activity. (B) Reactivated iSLK-WT cells were transfected with the K8.1 promoter reporter plasmid with or without the KSHV left lytic origin of replication cloned in *cis* and assessed for luciferase activity. (C) Reactivated iSLK-WT cells were transfected with the K8.1 and ORF57 promoter reporter plasmids in the presence or absence of DNA replication inhibitor (PAA) and assessed for luciferase activity. (D) Reactivated iSLK-WT and iSLK-ORF24_RAAAG_ mutant cells were transfected with K8.1 or ORF57 promoter reporter plasmids and assessed for luciferase activity. (E-F) iSLK-WT cells were transfected with plasmids containing the WT K8.1 promoter or the indicated single or double point mutant promoters and assessed for luciferase activity. The data for each mutant was normalized to the WT K8.1 promoter. All data shown are average of 3-5 biological replicates.

Notably, mutating the 5 bp RVNYS motif in the K8.1 promoter decreased transcription by > 50% (Figure 5E). This decrease was comparable to mutating the T at position four to an A in the TATTWAA sequence, which has been shown to be important (although not sufficient) for recognition and binding by ORF24 (10). Replacement of the 5 bp RVNYS motif with either synonymous (-S) or nonsynonymous (-NS) bases yielded a similar decrease in promoter activity (Figure 5E-K8.1_GGGAC->TACGA-S_, K8.1_GGGAC->ACTCG-NS_), suggesting a role for the overall architecture of this motif rather than the primary sequence. In support of this hypothesis, a preliminary analysis of four structural features of the expanded motif in the WT and mutant predict that the average roll of the DNA in the expanded motif is lower in the mutant promoter compared to the WT promoter (Figure S4). Combining the TATT->TATA mutation with the mutation of the 5 bp motif completely abolished transcription, indicating that these elements are absolutely essential for promoter activation (Figure 5E). Furthermore, with the exception of GG89TT-NS, systematic single and double base substitutions in the RVNYS motif were well tolerated, arguing that this region may not be contributing to base specific contacts with the vPIC (Figure 5F). In summary, while the expanded RVNYS motif appears important for transcriptional activation, we hypothesize it may contribute to vPIC specificity by impacting the local topology of the promoter DNA rather than through base specific contacts.

### The expanded promoter motif is important for DNA binding by vPIC

The RVNYS motif could either contribute to vPIC binding to its target promoters, as is the case for the TATTWAA sequence, or it could modulate downstream events involved in transcription initiation at vPIC bound promoters. To distinguish these possibilities, we used ChIP to measure HA-ORF34 occupancy on the luciferase plasmids containing WT or mutant K8.1 promoter in lytically reactivated HA-ORF34 iSLK cells. We engineered qPCR primers that enabled distinction of the K8.1 promoter on the plasmid DNA from the endogenous viral locus (Figure 6A).

**Fig 6:**
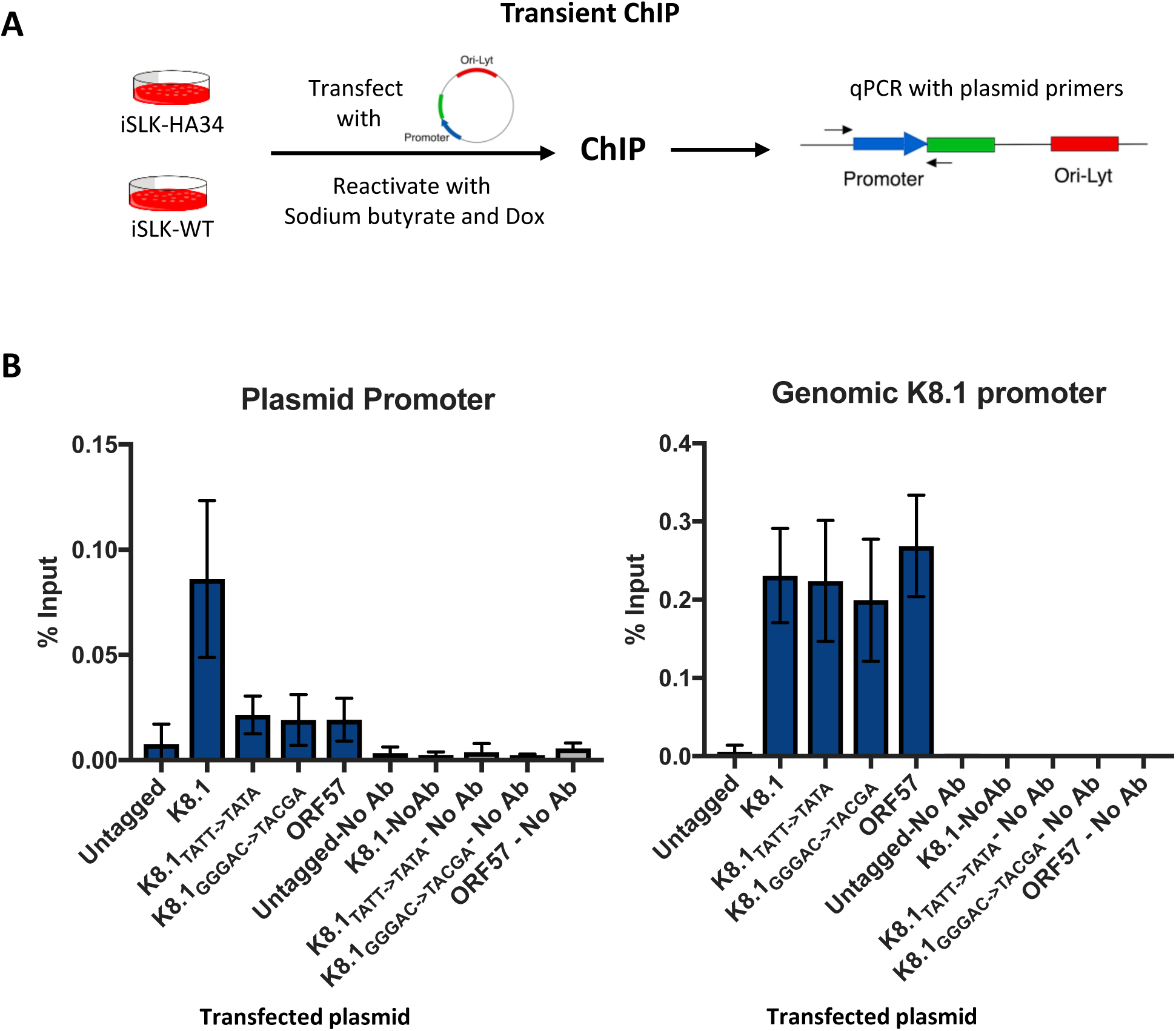
The expanded motif is important for DNA binding by vPIC. (A) Schematic of the transient ChIP assay. iSLK-WT or iSLK-HAORF34 cells reactivated with doxycycline and sodium butyrate were transfected with plasmid carrying the indicated promoter sequence. 48 h post transfection and reactivation, cells were subjected to ChIP with an anti-HA antibody, and the associated DNA was quantified using primers specific to the promoter on the plasmid and the genomic K8.1 promoter locus. (B) iSLK-HAORF34 cells were transfected with K8.1 promoter containing plasmid, mutant plasmids or ORF57 promoter plasmid (as a control). iSLK-WT cells without the HA tag transfected with K8.1 plasmid served as a control for the IP. The ChIP DNA was quantified using plasmid specific primers (left panel) and K8.1 promoter specific primers (right panel), normalized to the level of input DNA and presented as percent input. Data shown are an average of 3-6 biological replicates.

Notably, both the K8.1_TATT->TATA_ and K8.1_GGGAC->ACTCG_ mutations similarly impaired HA-ORF34 binding to the plasmid promoter DNA (Figure 6B-left panel). The ChIP signal was specific for HA-ORF34, as we observed only a background signal when the assay was performed in the WT iSLK cells containing an untagged ORF34 locus. Furthermore, there was comparable vPIC assembly at the genomic K8.1 locus in cells transfected with WT or the mutant plasmids, confirming similar immunoprecipitation efficiencies across samples (Figure 6B - right panel). Together, these data indicate that the expanded promoter motif is important for vPIC assembly on the promoter DNA.

### The ORF24-34 binding element in the origin is necessary for late gene activation

Given that late gene transcription requires the presence of the viral lytic origin of replication (Figure 5B), binding of ORF34 and ORF24 at the KSHV Ori was particularly notable (see Figures 3A-B). The binding site mapped to a region of the Ori just outside of the core minimal region that is required for viral DNA replication (Figure 7A). To examine the role of vPIC Ori binding on K8.1 promoter activation, we engineered a mutant version of the K8.1 promoter-driven luciferase plasmid lacking the ORF24-34 binding element (BE). A deletion encompassing this region has been shown to only modestly reduce (~30%) viral DNA replication (35,36). However, we observed a marked reduction in K8.1 promoter activation in the absence of the ORF24-34 Ori binding element (Figure 7B). The element by itself was insufficient to activate transcription in the absence of core minimal Ori, in agreement with the requirement for DNA replication to license late gene activation.

**Fig 7:**
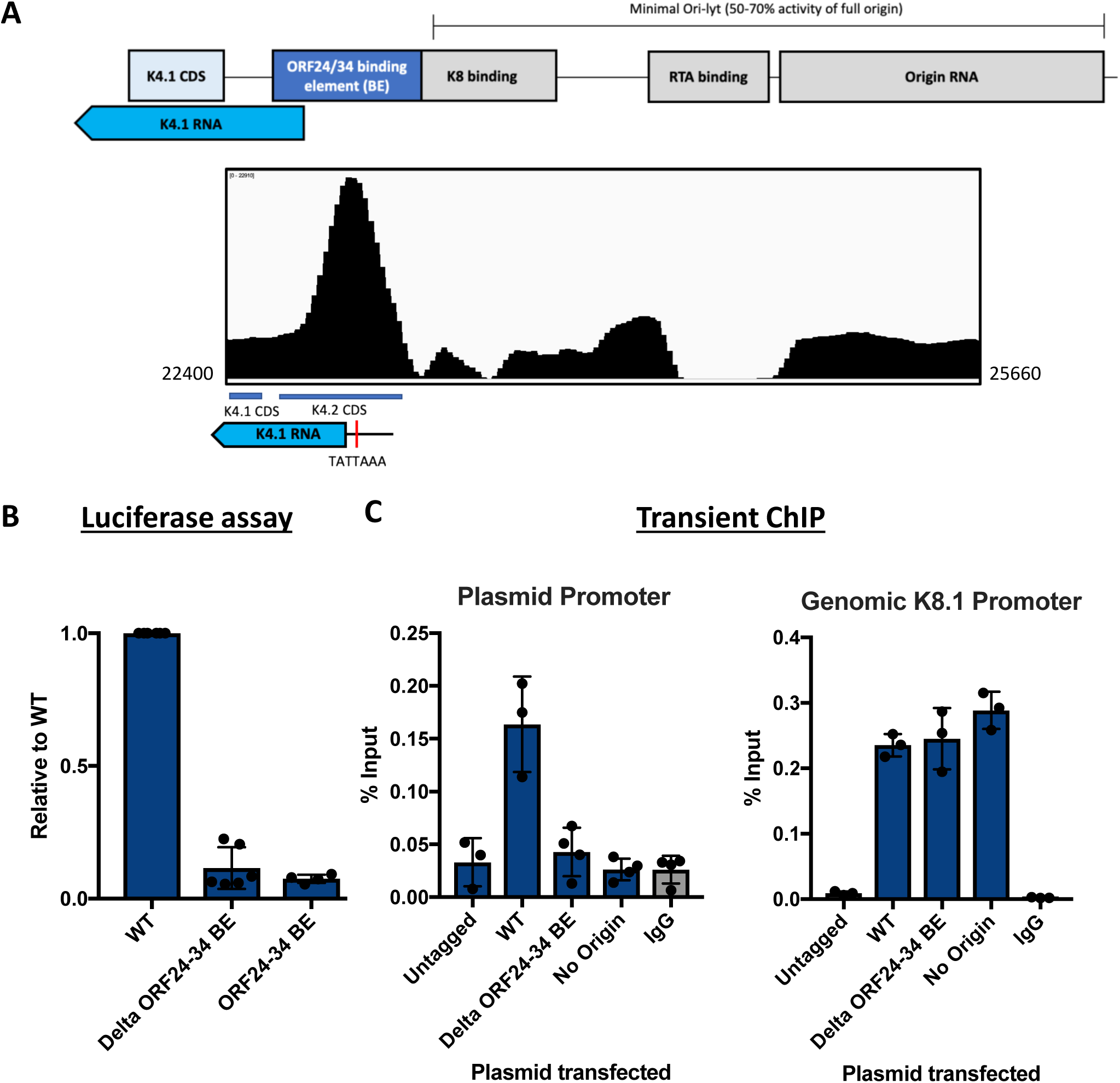
The ORF24-34 binding element at the origin is essential for late gene activation. (A) (Top panel) Schematic of the left lytic origin of replication of the KSHV genome. (Bottom panel) IGV image of the ChIP peak for HA-ORF34 at the origin of replication. (B) iSLK-WT cells were transfected with the K8.1 promoter plasmid containing either the full length Ori (WT), the Ori lacking the ORF24-34 binding element (Delta ORF24-34 BE), or the Ori lacking the minimal Ori-lyt and retaining only the ORF24-34 binding element (ORF24-34 BE). 48 h post transfection and reactivation, the cells were assessed for luciferase activity. (C) iSLK-HAORF34 cells were transfected with a K8.1 promoter containing plasmid containing either the full-length Ori (WT), the Ori lacking the ORF24-34 binding element (Delta ORF24-34 BE), or no Ori. Untagged iSLK-WT cells transfected with K8.1 plasmid served as a control for the IP. The ChIP DNA was quantified using plasmid specific primers (left panel) and genomic K8.1 promoter specific primers (right panel), normalized to the level of input DNA and presented as percent input. Data shown are an average of 3-4 biological replicates.

Ori binding by ORF24 and ORF34 could impact late gene transcription either by enabling vPIC promoter binding or by facilitating a downstream stage of transcription. To distinguish between these possibilities, we used our transient ChIP assay to compare HA-ORF34 binding to the K8.1 promoter on the plasmid containing either the WT Ori or the Ori lacking the ORF24-34 binding element. Removing the binding element within the Ori resulted in near complete loss of HA-ORF34 binding to the K8.1 promoter on the plasmid (Figure 7C), indicating that ORF24-34 occupancy at the lytic origin of replication is essential for vPIC assembly at the late gene promoter.

## Discussion

Late gene expression is uniquely regulated in KSHV by a set of viral proteins collectively known as the vPIC. In this study, we generated a comprehensive list of direct targets of the vPIC by combining transcriptome and genome-wide binding data of the complex. Among the direct targets, we identified an expanded promoter recognition motif that contributes to stable binding of the vPIC on the promoter. This is particularly useful to understanding alternate mechanisms of transcription initiation by RNA pol II because of the unique ability of the vPIC to directly recruit RNA Pol II to highly compact promoters. We also identified a vPIC binding element in the lytic origin of replication that licenses late gene activation at a distal promoter, suggesting a novel mechanism may underlie the link between DNA replication and late gene transcription.

Several key points emerge from independent analysis of the RNA Seq and ChIP Seq data. First, the previously identified TATTWAA motif is insufficient to mark binding and regulation by the vPIC, as demonstrated by the multiple ORFs that have the motif in their promoter (23/83 ORFs) but are not repressed in the ORF24_RAAAG_ mutant (10/23 ORFs) or bound by ORF34 (7/23 ORFs). Second, the TATTWAA motif is not a pre-requisite for regulation or binding by the vPIC. For example, although ORF53 expression is dependent on the ORF24-Pol II interaction and its promoter is bound by ORF34, its promoter does not have the conserved TATTWAA sequence but a TATATAA sequence in its place. Third, several genes that exhibit late kinetics escape repression in the mutant, indicating that not all late genes are direct or indirect targets of vPIC. It is notable that several of the kinetic late genes that escape repression are involved in immune regulation, consistent with a similar observation in EBV (28). In particular, EBV BPLF1 is a late gene whose expression is independent of the vPIC, and we observe that its homolog ORF64 also shows a similar pattern in KSHV. Finally, while there are several early genes that are indirect targets of the vPIC, only 1 of 13 direct targets exhibited early kinetics (ORF45). It is notable that ORF45 is a tegument protein, which might explain the requirement for its continued regulation at later times of infection.

Our integrative analysis revealed that a distinct 5 bp motif is integral to vPIC promoter recognition. Rather than engaging in base specific contacts (as is the case for the TATTWAA sequence) (7), we instead hypothesize that the importance of this motif lies in its shape or structure. In this regard, a preliminary analysis of basic structural properties of the expanded motif suggest that the average roll (representing the rotational flexibility) of the DNA in the expanded motif is lower in the 5 bp mutant promoter compared to WT K8.1 promoter. Structure-based recognition and a contribution of base pairs flanking the core binding site to binding affinity is a common feature among transcription factors, although the specific structural properties that influence binding differ for various classes of factors (37). A previous study of the bZIP family of transcription factors identified a correlation between the roll of DNA sequences that flank the core recognition element and binding affinity (37). Consistent with that, we observed a defect in the assembly of the vPIC when the expanded motif was mutated, suggesting a stabilizing role for the flanking sequence.

The location of the expanded motif is reminiscent of the downstream recognition element of the general transcription factor TFIIB (BRE^d^) (38). Interestingly, the BRE^d^ element also shows low sequence conservation and does not require an active promoter sequence to conform with the consensus sequence at all positions (38), as seen with the expanded motif. Interaction of TFIIB with the BRE^d^ element is thought to stabilize the TFIIB-TBP interaction on the promoter DNA, leading to activation or repression of the promoter in a context-dependent manner. While there is some evidence to suggest that TFIIB is present at late gene promoters (7), whether the interaction between the vPIC and the expanded motif is mediated by TFIIB or any other factors remains to be elucidated.

Notably, we found that KSHV DNA replication was reduced in the absence of functional ORF24, including with an ORF24 null mutant or ORF24 point mutants that fail to bind either RNA Pol II or the vPIC component ORF34. This was surprising, given that deletion or mutation of several other vPIC components has been shown not to affect DNA replication (7,13,17,18). Furthermore, we previously failed to observe a replication defect in an ORF24 null virus (7). We hypothesize that the discrepancy for the ORF24 null mutant relates to the methods used to establish the infected cell lines. In our earlier study, the viral BAC DNA was directly transfected into iSLK cells, resulting in only a ~5-7-fold increase in genome copy number upon reactivation. It is likely that the majority of viral DNA was present in a non-reactivatable state, providing a level of background that significantly reduced the dynamic range for DNA replication measurements. Here, the iSLK cells were established through co-culture with reactivated 293T cells harboring the BAC DNA, i.e. through infection rather than transfection. This reproducibly yields ~50-70-fold increases in genome copy number upon lytic reactivation, more closely representing actual infection. That said, the role of ORF24 in viral DNA replication may be unique to KSHV, as this link has not been observed upon depletion of its homologs in EBV and MHV68 (28,39).

One possibility for how KSHV ORF24 might contribute to DNA replication is through its indirect transcriptional targets. We observed that components of the helicase/primase complex (ORFs 40, 41), which are involved in DNA replication, show more than average repression in the absence of functional ORF24 (Figure 2). Importantly, these genes were not repressed in previous studies using stop mutants of the vPIC components ORF18, ORF30 and ORF31 (17,18), raising the possibility that ORF24 has additional gene regulatory functions outside the context of the vPIC. For example, our observed binding of ORF24 and ORF34 at the lytic origins of replication could facilitate DNA replication. Consistent with this hypothesis, two previous studies that defined the boundaries of the lytic origin of replication observed a modest decrease in replicated DNA upon deletion of the ORF24-34 binding site identified here (35,36).

Interestingly, we also observed that binding of ORF24-34 at the origin is critical for origin dependent late gene expression, as deletion of the ORF24-34 binding element at the origin prevented vPIC assembly and transcription at the distal K8.1 late promoter. We note that although the binding element overlaps the promoter of K4.1, we observe less than average reduction in K4.1 transcript levels in the mutant compared to the mutant rescue. We hypothesize that alternate mechanisms of maintaining steady-state K4.1 RNA levels are adopted in the absence of a functional vPIC and binding of ORF24-34 at the origin is likely related to the link between DNA replication and late gene activation. The proximity of the ORF24-34 binding element to the other origin elements also raises the possibility of vPIC recruitment by other origin binding proteins. Exploring the mechanism underlying this link between origin binding and late gene activation will be an exciting area for future study.

## Methods

### Cell lines

HEK293T cells were maintained in DMEM (Invitrogen) + 10% Fetal Bovine Serum (FBS). iSLK BAC16 cells (29) were maintained in DMEM + 10% FBS and 0.5 mg/mL hygromycin B. HEK293T cells stably expressing ORF24-3xFlag were maintained in DMEM (Invitrogen) + 10% Fetal Bovine Serum (FBS) and 325 ug/ml of zeocin. iSLK-puro cells were maintained in DMEM + 10% FBS.

### Plasmids

A 100 bp fragment of the K8.1 promoter or ORF57 promoter was generated by PCR from the BAC16 genome and cloned into the KpnI and HindIII sites of the pGL4.16 reporter plasmid using Infusion cloning (Clontech). The left origin of replication was similarly generated by PCR from the BAC16 genome and cloned into the NotI and BstXI sites of the K8.1Pr-pGL4.16 plasmid. The promoter mutants were generated by site-directed mutagenesis using KAPA HiFi DNA polymerase using the primers shown in Supplementary table 1. To make the ORF24-3xFlag pJLM1 plasmid for lentiviral transduction, the ORF24-3xFlag fragment was generated by PCR using ORF24-3xFlag-pCDNA4 as the template and cloned in to pLJM1 plasmid digested with AgeI and EcoRI. All primers used for cloning and mutagenesis are listed in Supplementary table 1.

### Generation of BAC mutants and establishing latent cell lines

The ORF24_RAAAG_, ORF24_R328A_ and ORF24_Stop_ mutants, the corresponding mutant rescues (MR) for ORF24_RAAAG_, ORF24_R328A_ and N terminal HA tagged ORF24 and ORF34 were engineered using the scarless Red recombination system in BAC16 GS1783 *E. coli* (40) as described previously. The modified BACs were purified using the NucleoBond BAC 100 kit (Clontech).

To establish iSLK cell lines latently infected with the modified virus, HEK293T cells were transfected with 5-10 ug of the modified BAC16 using linear polyethylenimine (PEI, MW ~25,000) at a 1:3 DNA:PEI ratio. The following day, the 293T cells were mixed 1:1 with iSLK-puro cells (which contain inducible RTA but lack KSHV) and treated with 30 nM 12-O-Tetradecanoylphorbol-13-acetate (TPA) and 300 nM sodium butyrate for 4 days to induce lytic replication. Cells were then grown in selection media containing 300 ug/ml hygromycin B, 1 ug/ml puromycin and 250 ug/ml G418. The hygromycin concentration was gradually increased to 500 ug/ml and then 1 mg/ ml until all the HEK293Ts cells died. To generate the ORF24_RAAAG_, ORF24_R328A_, and ORF24_Stop_ (which are defective in infectious virion production), HEK293T cells that stably express ORF24-3xFlag were used for the initial transfection and co-culture.

### Supernatant transfer assay

For supernatant transfer assays to assess virion production, BAC16-containing iSLK cells were plated at a density of 1×10^6^ cells in 10 cm dishes and treated with 1 ug/mL doxycycline and 1 mM sodium butyrate the following day. 72 h post induction, the supernatant was filtered through a 0.45 uM syringe filter and 2 ml of the supernatant was mixed with 1×10^6^ freshly trypsinized HEK293T cells in a 6 well plate and centrifuged at 1,500x*g* for 2 h at 37°C. The following day, cells were trypsinized, fixed in 4% paraformaldehyde and the percentage of cells expressing GFP was determined by flow cytometry (BD Csampler, BD Biosciences).

### Viral genome replication

To measure viral genome replication, iSLK-BAC16 cells were reactivated for 72 h as described above. The cells were then scraped into the media and the media + cells were digested with proteinase K (80μg/mL) (Promega) in 5x proteinase K digestion buffer (50mM Tris-HCl pH 7.4, 500mM NaCl, 5mM EDTA, 2.5% SDS) overnight at 55°C. The gDNA was isolated using Zymo Quick gDNA Miniprep Kit according to the manufacturer’s instructions. Quantitative PCR (qPCR) was performed on the isolated DNA using iTaq Universal SYBR Green Supermix on a QuantStudio3 Real-Time PCR machine. DNA levels were quantified using relative standard curves with primers specific for KSHV ORF59 promoter and human CPSF6 promoter (Supplementary table 1). The relative genome numbers were normalized to CPSF6 to account for loading differences and to uninduced samples to account for differences in starting genome copy number.

### RT-qPCR

To look at viral transcript levels, total RNA was isolated from reactivated iSLK cells at the indicated time points using the Zymo Direct-Zol RNA kit following the manufacturer’s protocol. The purified RNA was treated with DNase and cDNAs were synthesized using AMV reverse transcriptase (Promega). The cDNAs were directly used for qPCR analysis using the iTaq Universal SYBR Green Supermix. The qPCR signal for each ORF was normalized to 18s rRNA.

### Western blotting

Cells were lysed in 1× lysis buffer (50mM Tris-HCl pH 7.5, 150mM NaCl, 0.5% NP-40, cOmplete EDTA-free Protease Inhibitors [Roche]) by rotating for 30 min at 4°C. The lysate was clarified by centrifugation at 21,000 x *g* for 15 min at 4°C. The supernatant was quantified by Bradford assay, and equivalent amounts of each sample were resolved by SDS-PAGE and western blotted with rabbit polyclonal antibodies against vinculin (Abcam, 1:1000), ORF59, or K8.1. The ORF59 and K8.1 antibodies were generated by injecting rabbits with purified MBP-ORF59 or MBP-K8.1 (gifts from Denise Whitby (41)) (Pocono Rabbit Farm and Laboratory).

### RNA Seq sample preparation and analysis

Total RNA was isolated from reactivated iSLK cells at indicated time points using Zymo Direct-Zol RNA kit following the manufacturer’s protocol. The samples were ribo-depleted and stranded libraries were prepped by the QB3 Berkeley core sequencing facility. Multiplex sequencing was performed using the Illumina HiSeq 4000 to generate 100 bp paired end reads. The quality of the raw reads was checked using FastQC (42) and adapters were trimmed using TrimGalore (43) using default parameters. The trimmed reads were aligned to both the viral (NC_009333.1) and host genomes (hg19) using STAR aligner (ver2.5.3a) (44). The aligned reads were counted using htseq-counts (45) and differential expression was assessed using DESeq2 (46).

To compensate for the problem of quantitative assessment of transcripts due to the overlapping transcripts in the KSHV genome, the primary counts of the transcripts were calculated as recommended in Bruce et al (30) with the following small modifications. Length correction was not applied to allow for differential expression analysis by DESeq2. Transcripts for which the counts became negative in all 3 replicates when calculating primary counts were removed before running DESeq2. Bar plots showing Log_2_ Fold Change were generated using R. The heatmap was generated using the heatmap.2 package in R. The fold change values were scaled by column before plotting to ease visualization of the data. Raw and processed data files are available from the GEO database (accession number: GSE126602).

### Chromatin Immunoprecipitation (ChIP)

ChIP was performed on 3*15-cm plates of iSLK-cells (WT KSHV, KSHV_HA-ORF34_ and KSHV_ORF24-HA_) reactivated for 48 h with 1 ug/ml dox and 1 mM sodium butyrate. Cells were crosslinked in 2% formaldehyde for 15 min at room temperature, quenched in 0.125 M glycine, and washed twice with ice-cold PBS. Crosslinked cell pellets were mixed with 1 ml ice-cold ChIP lysis buffer (5mM PIPES pH 8.0, 85mM KCl, 0.5% NP-40) and incubated on ice for 10 min, whereupon the lysate was dounce homogenized to release nuclei and spun at 1700 g for 5 min at 4°C. Nuclei were then resuspended in 800 μl of nuclei lysis buffer (50mM Tris-HCl pH 8.0, 0.3% SDS, 10mM EDTA) and rotated for 10 min at 4°C followed by sonication using a QSonica Ultrasonicator with a cup horn set to 25 amps for 20 min total (5 min on, 5 min off). Chromatin was spun at 16,000 x *g* for 10 min at 4°C and the pellet was discarded. The chromatin was precleared with protein A + protein G beads blocked with 5 ug/ml glycogen, 50 ug/ml BSA, 100 ug/ml Ecoli tRNA in 1ml dilution buffer (1.1% Triton-X-100, 1.2mM EDTA, 16.7mM Tris-HCl pH 8, 167mM NaCl) for 2 hours.

350 μl of chromatin was diluted in ChIP dilution buffer (16.7 mM Tris-HCl pH 8.0, 1.1% Triton X-100, 1.2 mM EDTA, 167 mM NaCl) to 500 μl and incubated with 10 μg anti-HA antibody (Cell Signaling C29F4) overnight, whereupon samples were rotated with 20 μl pre-blocked protein A + G beads (Thermofisher) for 2 h at 4°C. Beads were washed with low salt immune complex (20 mM Tris pH 8.0, 1% Triton-x-100, 2 mM EDTA, 150 mM NaCl, 0.1% SDS), high salt immune complex (20 mM Tris pH 8.0, 1% Triton-x-100, 2 mM EDTA, 500 mM NaCl, 0.1% SDS), lithium chloride immune complex (10mM Tris pH 8.0, 0.25 M LiCl, 1% NP-40, 1% Deoxycholic acid, 1 mM EDTA), and Tris-EDTA for 10 min each at 4°C with rotation. DNA was eluted from the beads using 100 μl of elution buffer (150 mM NaCl, 50 μg/ml Proteinase K) and incubated at 50 °C for 2 h, then 65 °C overnight. DNA was purified using the Zymo PCR purification kit. For ChIP-qPCR, purified DNA was quantified by qPCR using iTaq Universal SYBR Mastermix (BioRad) and the indicated primers (Supplementary table 1). Each sample was normalized to its own input.

For ChIP-Seq, the concentration of the eluted DNA was measured using Qubit. 10-15 ng of eluted DNA was used to generate libraries using the Accel-NGS 2S Plus DNA Library Kit (Swift Biosciences). The libraries were sequenced on a Illumina HiSeq 4000 sequencer at the QB3 Berkeley core facility to generate 100 bp single end reads.

### ChIP Seq Analysis

The quality of the raw reads was checked using FastQC (42) and adapters were trimmed using TrimGalore (43) using default parameters. The trimmed reads were aligned to both the viral and host genomes (hg19) using Bowtie2 aligner using default parameters. The aligned reads were filtered for uniquely mapped reads using SAMTools. The filtered reads were converted to tdf format using IGV Tools and visualized on Integrated Genome Viewer (47,48). We used HOMER (33) with the following minor adjustments to default parameters to call peaks for the immunoprecipitated and input samples from cells that have HA tagged ORF34 and untagged ORF34 (WT) as a control. The fold over input and local fold change were set to 1.3 as recommended for smaller and denser viral genomes (personal communication with Chris Benner, UCSD). The peak size and fragment length were also set at 200 and 150 respectively. From the resulting peaks, only the ones that were present in both conditions (HA-ORF34 vs HA-Input, HA-ORF34 vs WT) and both replicates were retained for further analysis. The common peaks identified were verified visually on the Integrated Genome Viewer and were manually annotated.

To identify the direct targets, transcripts that had a Log_2_ fold change lower than the mean Log_2_ fold change in the ORF24_RAAAG_ mutant vs MR were considered to be down regulated in the mutant. Although ORF8 and ORF26 were not included in the differential expression analysis, they were considered to be downregulated because of negative values for their levels in the mutant and not in the mutant rescue on calculation of primary transcript levels. Raw and processed data files are available from the GEO database (accession number: GSE126601).

### Promoter Analyses

We defined the promoter sequence as 100 base pairs (bp) upstream from the transcription start site (TSS) or 300 bp upstream from the translation start site when the TSS was unknown. The TSS data was obtained from Table 1 in (23) and a custom python script was used to extract the sequences. These were then run through MEME (31) allowing for multiple occurrences of a motif in a sequence and only searching the given strand to identify enriched motifs. The identified motif was then used to search for similar motifs across all 83 promoter sequences from the viral genome, again only allowing for searching the given strand.

### Reporter assays

5*10^5^ iSLK-BAC16 (WT or ORF24_RAAAG_) cells were plated in a 6 well plate 24 h before transfection and reactivation. The cells were reactivated with 1 μg/ml doxycycline and 1 mM Sodium Butyrate and simultaneously transfected with 0.95 μg luciferase reporter plasmid and 0.05 μg renilla plasmid using polyjet transfection reagent. 48 h post transfection and reactivation, the cells were rinsed twice with room temperature PBS, lysed by rocking for 15 min at room temperature in 300 μl of Passive Lysis Buffer (Promega), and clarified by centrifuging at 14,000 g for 1 min. 20 μl of the clarified lysate was added in triplicate to a white chimney well microplate (Greiner bio-one) to measure luminescence on a Tecan M1000 using a Dual Luciferase Assay Kit (Promega). The firefly luminescence was normalized to the internal Renilla luciferase control. The induced sample was then normalized to the uninduced (but transfected) control to look at the fold induction during reactivation.

### Transient ChIP-qPCR

3.5×10^6^ iSLK-BAC16 were plated in a 15 cm dish 24 h before transfection and reactivation. The cells were reactivated with 1 μg/ml doxycycline and 1 mM Sodium Butyrate and simultaneously transfected with 8 μg of reporter plasmid using polyjet transfection reagent. 48 h post reactivation and transfection, cells were crosslinked and lysed as mentioned above. Nuclei were sonicated using a Covaris S220 (Covaris) for 5 min with the recommended sonication settings (Power: 140 W, Bursts per cycle: 200, Duty cycle: 5%). The chromatin was pre-cleared as mentioned above and the concentration measured using a qubit. 25 μg of pre-cleared chromatin was used in the IP and the following steps were same as mentioned in the ChIP protocol above.

## Acknowledgements

We thank all members of the Glaunsinger and Coscoy labs for their helpful suggestions. In particular, we thank Angelica Casteñeda for providing the HEK293T cells stably expressing ORF24-3xFlag and Chris Benner (UCSD) for suggestions on proper usage of HOMER for organisms with small genomes. This work was supported by NIH R01 AI122528 to B.G., who is also a Howard Hughes Medical Institute Investigator.

---

**Fig S1:**
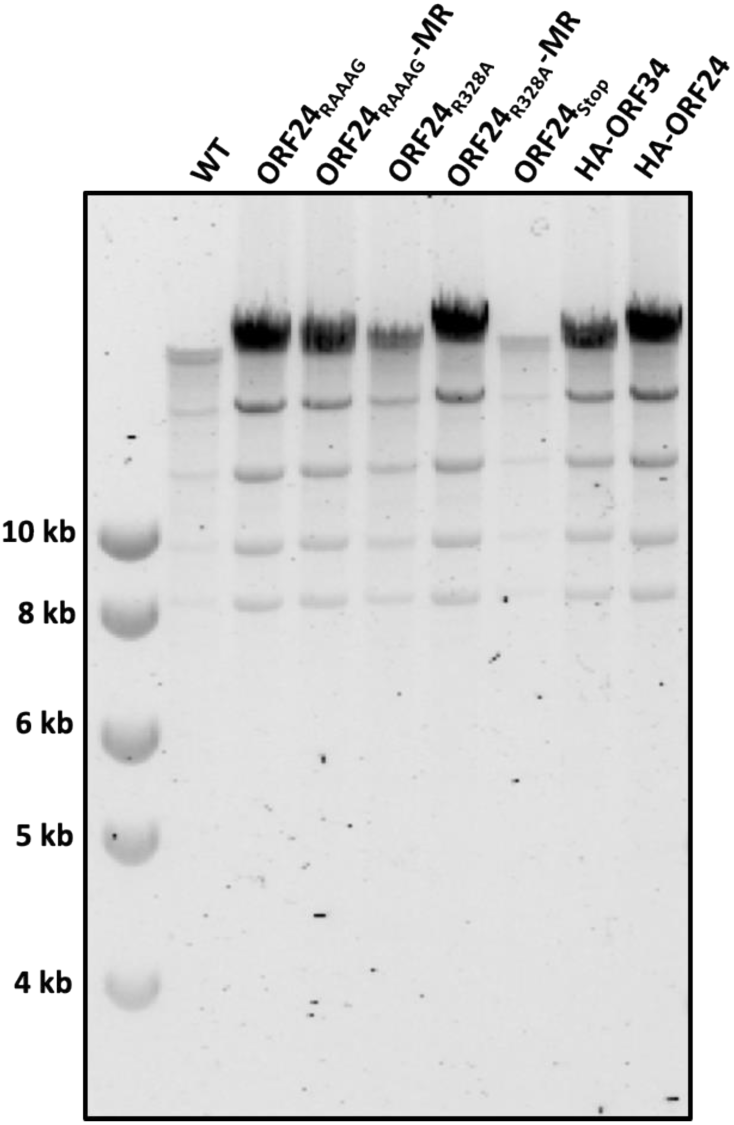
Validation of mutant BAC DNA integrity. The integrity of the various BAC mutants used in this study was verified by *Rsr*II digestion.

**Fig S2:**
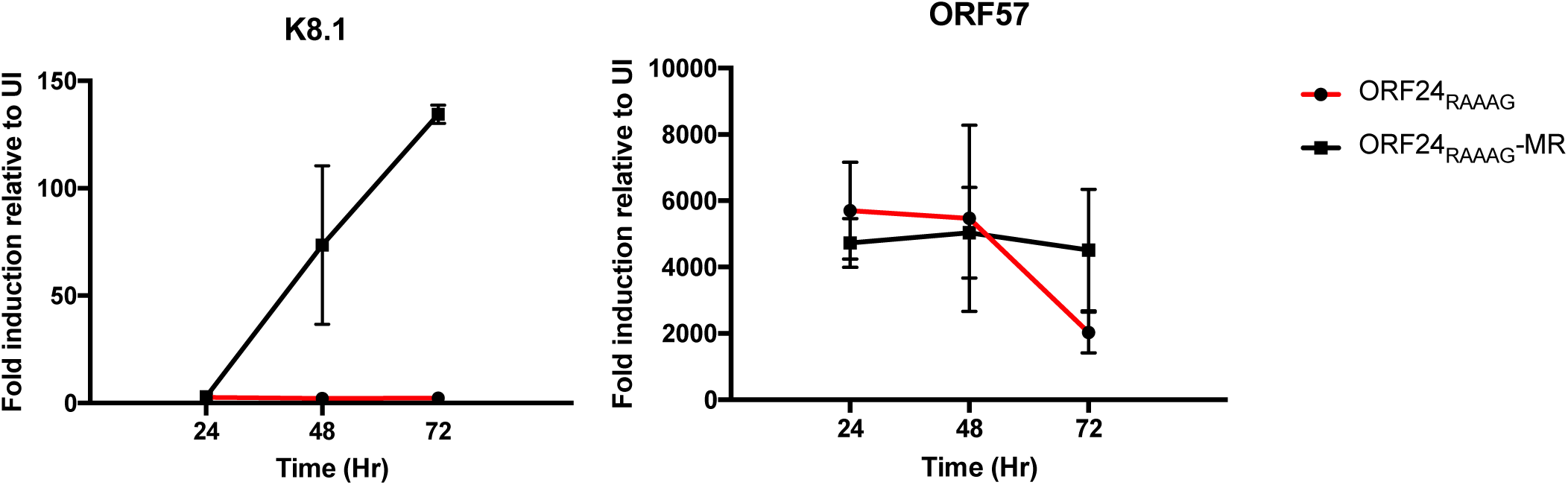
Kinetics of early and late gene transcripts in ORF24_RAAAG_ mutant and MR. iSLK-ORF24_RAAAG_ and MR were reactivated with doxycycline and sodium butyrate for the indicated time points, whereupon total RNA was extracted, converted to cDNA, and quantified by qPCR using primers specific to the CDS of K8.1 (late gene) or ORF57 (early gene). The fold induction was measured relative to uninduced sample.

**Fig S3:**
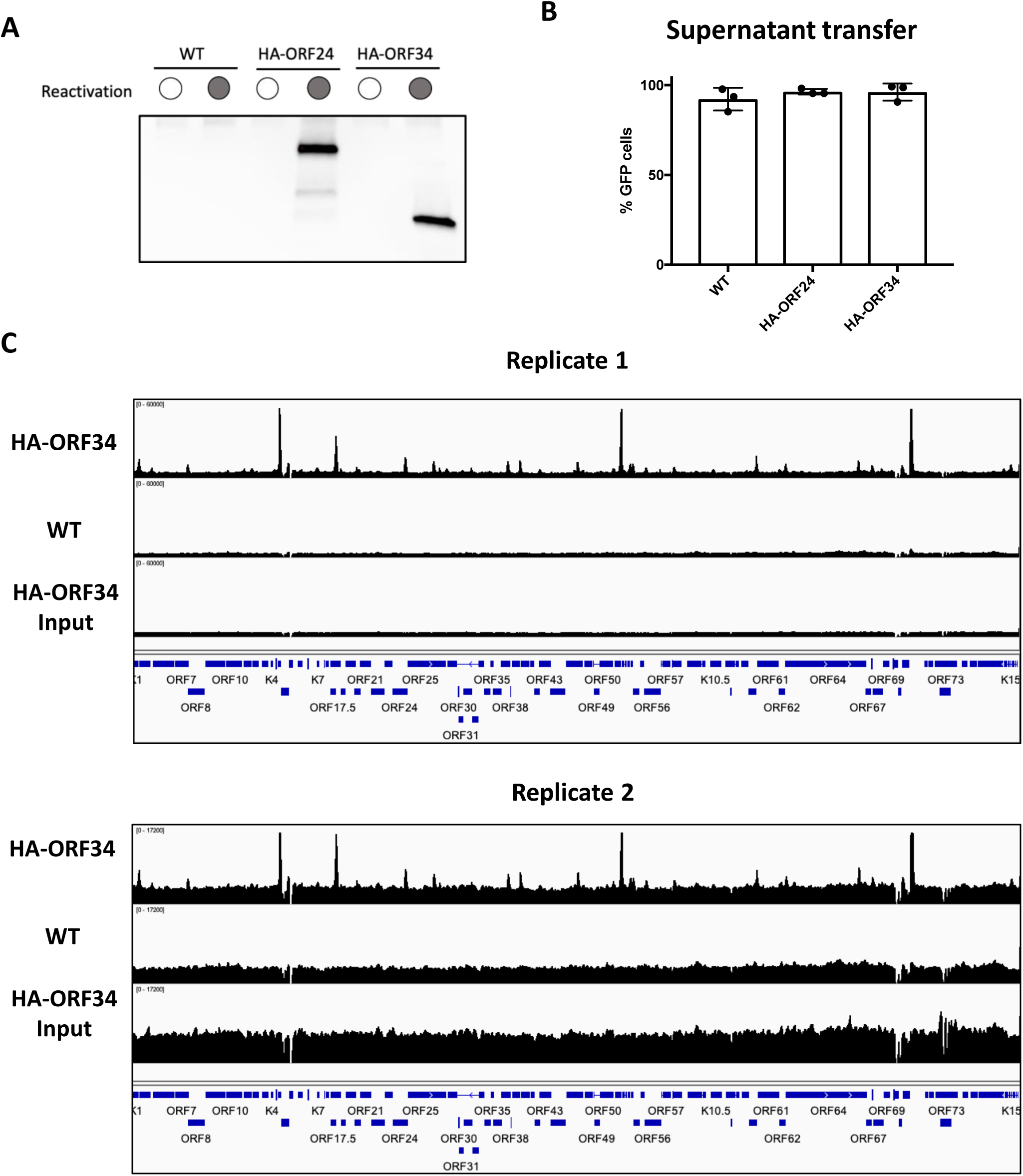
ChIP-Seq data with HA-ORF34. (A) Western blot showing expression of HA-ORF24 and HA-ORF34. iSLK cells harboring WT BAC16, HA-ORF34 BAC or HA-ORF24 BAC were reactivated with doxycycline and sodium butyrate for 72 hours. Protein lysate was immunoprecipitated with HA beads and visualized by SDS-PAGE-Western blot. (B) iSLK cells harboring WT BAC16, HA-ORF34 BAC or HA-ORF24 BAC were reactivated with doxycycline and sodium butyrate for 72 hours. Progeny virion production was measured by supernatant transfer on to 293T cells and quantified by flow cytometry. (B) IGV image showing ChIP-Seq data from iSLK-HA-ORF34 cells reactivated with doxycycline and sodium butyrate for 48 h and immunoprecipitated with anti-HA antibody. Also shown are plots for control data from untagged iSLK cells and input DNA for two biological replicates

**Fig S4:**
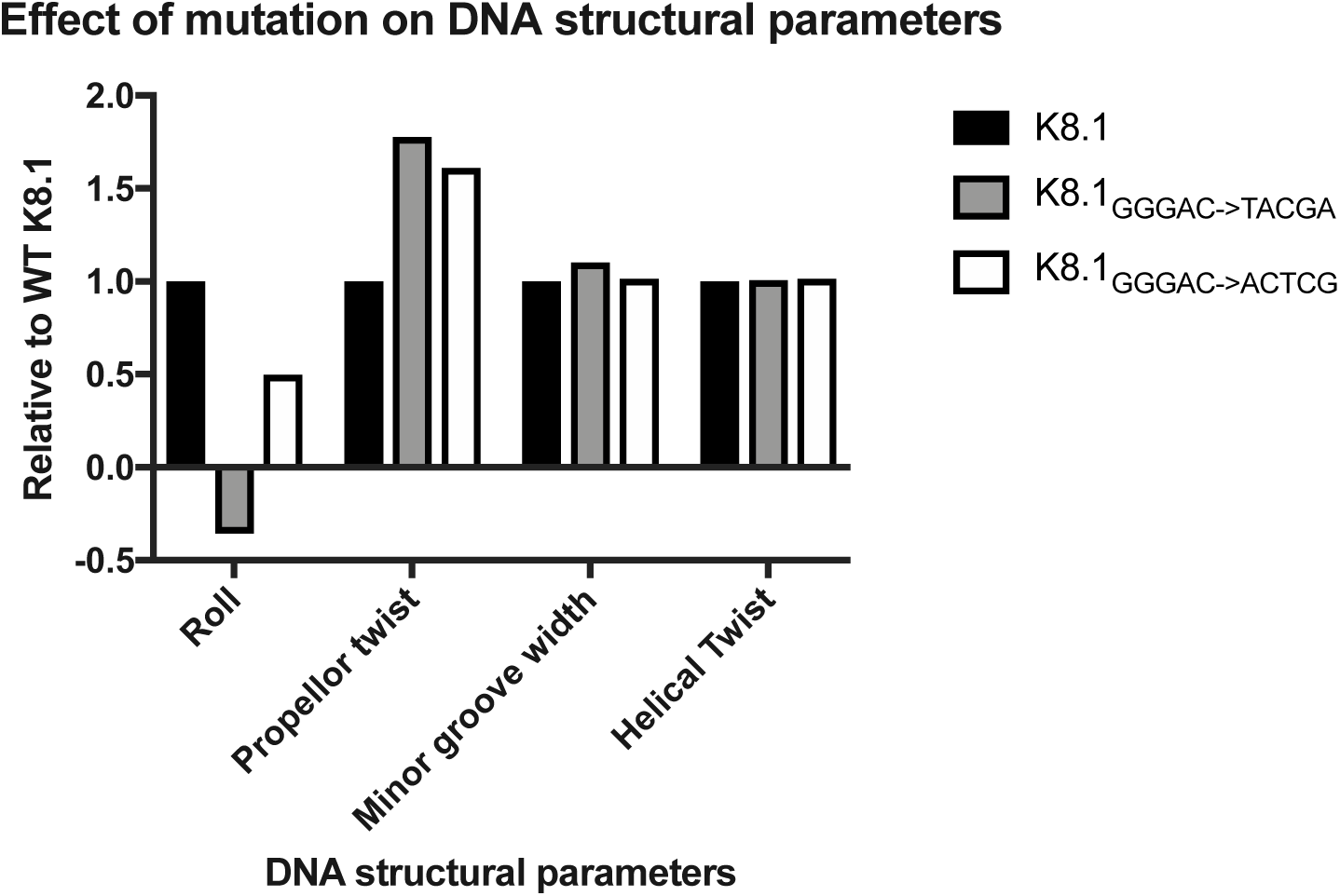
Effect of mutations on DNA structural parameters. Four structural parameters for 40 bp of the WT and expanded motif mutants were calculated using the DNAShape program in R (49). Average of each structural parameter for the mutated base pairs and the two adjacent base pairs at either end (a total of 7 bp) was calculated for each DNA sequence analyzed. The two adjacent base pairs were included in the averaging as some of the structural parameters refer to a base step rather than a specific base. The average for each mutant was normalized to the WT K8.1 promoter.

**Supplementary Table 1:**
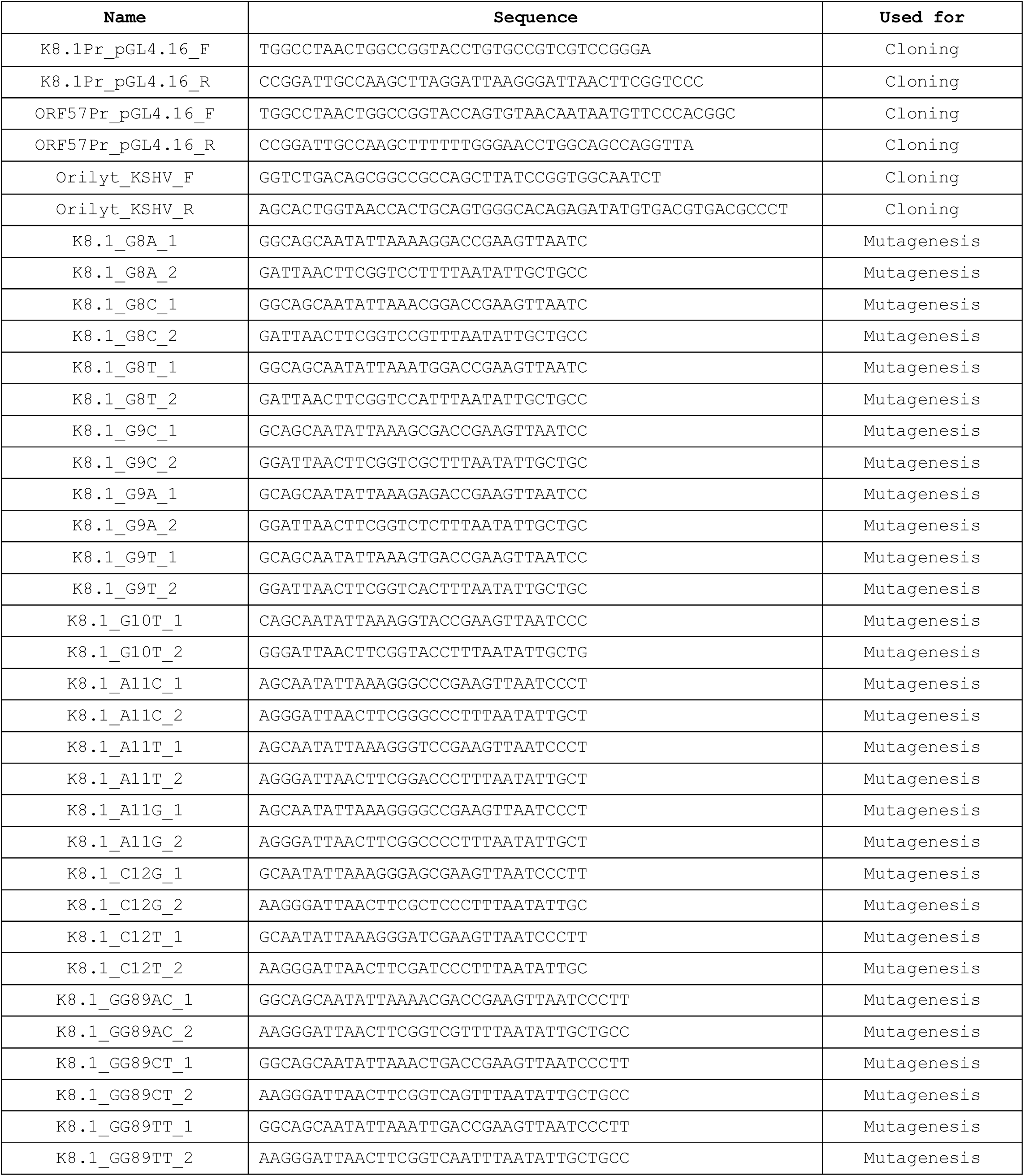
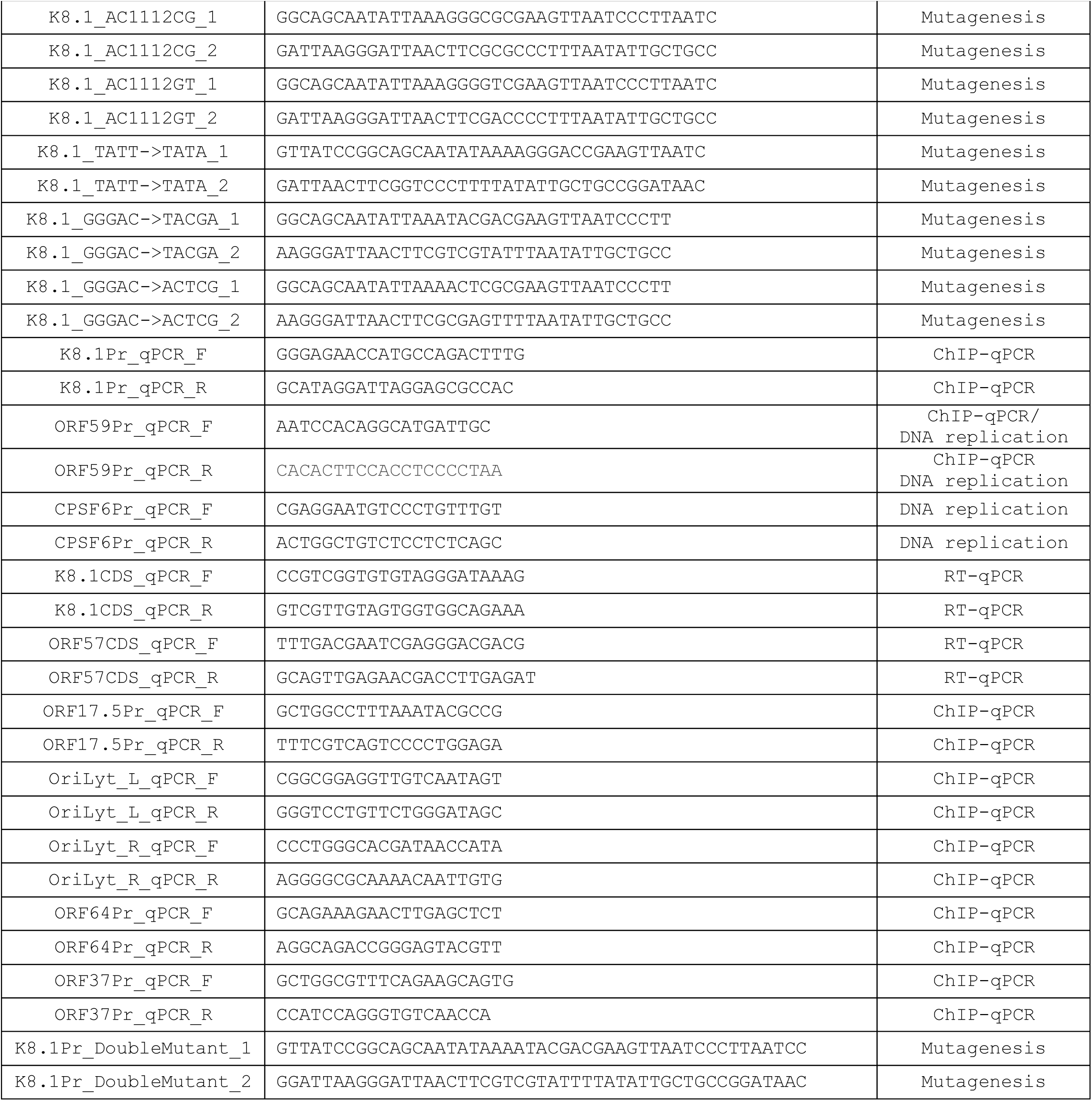
List of all DNA sequences used in the study

